# On the effectiveness of small, discriminatively pre-trained language representation models for biomedical text mining

**DOI:** 10.1101/2020.05.20.107003

**Authors:** Ibrahim Burak Ozyurt

## Abstract

Neural language representation models such as BERT [1] have recently shown state of the art performance in downstream NLP tasks and bio-medical domain adaptation of BERT (Bio-BERT [2]) has shown same behavior on biomedical text mining tasks. However, due to their large model size and resulting increased computational need, practical application of models such as BERT is challenging making smaller models with comparable performance desirable for real word applications. Recently, a new language transformers based language representation model named ELECTRA [3] is introduced, that makes efficient usage of training data in a generative-discriminative neural model setting that shows performance gains over BERT. These gains are especially impressive for smaller models. Here, we introduce a small ELECTRA based model named Bio-ELECTRA that is eight times smaller than BERT BASE and achieves comparable performance on biomedical question answering and yes/no question answer classification tasks. The model is pre-trained from scratch on PubMed abstracts using a consumer grade GPU with only 8GB memory. For biomedical named entity recognition, however, large BERT Base model outperforms both Bio-ELECTRA and ELECTRA-Small++.

## I. Introduction

Transformers based language representation learning methods such as Bidirectional Encoder Representations from Transformers (BERT) [1] are becoming increasingly popular for downstream biomedical NLP tasks due to their performance advantages [2]. The performance of these models comes at a steep increase in computation cost both at training and inference time, For example, we use a BERT based re-ranker as the final step in our biomedical question answering system Bio-AnswerFinder [4] (https://github.com/SciCrunch/bio-answerfinder), where 60% of the question answering time latency is due to the BERT classifier with 110 million parameters. The increased size of the transformer models is correlated with the increased performance [1]. Since the computational cost involved at inference time for large models is a bottleneck in their practical applications in the real world, new approaches to achieve similar performance on smaller models are getting increasingly popular. A popular approach on this end is distilling BERT to a smaller classifier such as DistillBERT [5], TinyBERT [6] and MobileBERT [7]. However, a small,efficient model without going through the trouble of training a large model and mimicking it in a smaller model is more preferable.

BERT uses a masked language modeling (MLM) approach by masking 15% of the training sentences and learning to guess the masked tokens in a generative manner. This results BERT using only 15% of training data. A recent approach called ELECTRA [3], introduced a new language modeling approach where a discriminative model is trained to detect whether each token in the corrupted input was replaced by a co-trained generator model sample or not. ELECTRA is computationally more efficient than BERT and outperforms BERT given the same model size, data and computation resources. The improvements over BERT is most impressive at small model sizes, which makes it an excellent candidate in pursuit of small and efficient language representation models for biomedical text mining.

In this paper, we introduce a small and efficient ELECTRA based domain-specific language representation model trained on PubMed abstracts with a domain specific vocabulary achieving comparable results on question answering related tasks to BERT Base model having 8 times more parameters resulting in 8 times decrease in inference time. The model is trained on a modest consumer grade GPU with only 8GB RAM which is much lower bar for pre-training of domain-specific language representation models that BERT and variants. The performance on biomedical named entity recognition of small ELECTRA models are not as impressive as in the question answering related tasks compared to BERT.

## II. Methods

### A. Pre-training Bio-ELECTRA

Both ELECTRA and BERT are pre-trained on English Wikipedia and BooksCorpus as general purpose language models.

Both BERT and ELECTRA use WordPiece tokenization [8] which represents words as constructed from character n-grams of highest co-occurrence to allow out-of-vocabulary (OOV) words to be represented. Given a vocabulary size, the character n-grams (subwords) making up the vocabulary are determined from the corpus by using an objective similar to the compression algorithms to find the subwords that would generate each unique word in the corpus. OOV words are generated by combination of subwords from the subwords vocabulary. Since the vocabulary of BERT and ELECTRA [3] are generated from general purpose corpora, a lot of biomedical domain specific words need to be composed from subwords that does not convey enough information by themselves. For example the gene BRCA1 in BERT/ELECTRA vocabulary represented as B##R##CA##1, mostly formed from single letter embedded representations. For Bio-ELECTRA, the vocabulary is generated using SentencePiece byte-pair-encoding (BPE) model [9] from PubMed abstract texts from 2017. Using this domain-specific vocabulary BRCA1 is represented as BRCA##1. In this case, the composition from parts conveys more information since the learned vector embedding of BRCA subword is more like to capture for example its breast cancer relatedness.

19.2 million most recent PubMed abstracts (having PMID greater than 10 million) as of March 2020 are used for Bio-ELECTRA pre-training. Sentences extracted from the paper title and abstract are used to build the pre-training corpus of about 2.5 billion words. Using the PubMed abstract corpus and 2017 PubMed abstracts generated SentencePiece vocabulary ELECTRA-Small model (14M trainable parameters) with a maximum sequence size of 256 and batch size of 64 is pre-trained from scratch on a RTX 2070 8GB GPU in four stages for 24 days for 1.8 million steps. Original ELECTRA Small was trained on a V100 32GB GPU in 4 days with a batch size of 128 for one million steps. However, the distributed ELECTRA Small++, which was used for our comparison experiments, was trained on the XLNet [10] corpus (about 33 billion subword corpus) with maximum sequence size of 512 for 4 million steps. Since the batch size of Bio-ELECTRA half the size of the ELECTRA Small due to our GPUs memory size, two million steps are equivalent to one million ELECTRA training steps. ELECTRA Small++ is trained four times more than Bio-ELECTRA and trained on much larger corpus.

### B. Fine-tuning for Biomedical Text Mining Tasks

The syntactic and semantic language modeling information latently captured in the pre-trained weights of transformer models combined with a classification layer were found to provide state-of-the-art results in many NLP tasks [1], [3]. We fine-tune Bio-ELECTRA, ELECTRA Small++ and BERT Base for biomedical question answering, yes/no question answer classification and named entity recognition (NER) tasks.

For biomedical question answering, we used BERT and ELECTRA architectures for SQuAD citesquad for SQuAD v1.1. Similar to Wiese et al and Lee et al. [2], [11], we have combined our BioASQ 8b training set generated factoid and list questions based training test with out-of-domain SQuAD v1.1 data set to increase performance over much smaller BioASQ data.

The biomedical yes/no question answer classification task is similar to sentiment (hedging for biomedical literature) detection where the polarity (positive/negative) of a candidate sentence needs to be detected in the context of a question. For ELECTRA and BERT, we have used their official codebase from GitHub slightly extended for our specific classification task.

Named entity recognition involves detection of names of biomedical entities in sentences and usually used for down-stream tasks such as information extraction and question answering. For ELECTRA and Bio-ELECTRA, we have used the ELECTRA architecture for entity level tasks adapted for BIO annotation scheme. For BERT, we have used HuggingFace Transformers library single output layer entity classification architecture.

For each fine-tuning experiment, ten randomly initialized models are trained and average testing performances and standard deviations are reported. Default BERT and ELECTRA hyperparameters including the number of epochs (two for QA task and three for classification/NER tasks) are used for corresponding experiments. More performance can be squeezed out of the fine-tuning models by hyperparameter tuning. However, since the main goal is comparing different models under similar conditions, this was not attempted. All of the ELECTRA based fine-tuning trainings are conducted on a GTX 1060 6GB GPU, while 8x larger BERT model required training on our RTX 2070 8GB GPU. For BERT experiments cased BERT Base is used.

## III. Results

### A. Datasets

For biomedical question answering and yes/no answer classification tests, we have generated training and testing data sets from the publicly available 2020 BioASQ [12] Task B (8b) training data set. BioASQ 8b training set consists of 3243 questions together with ideal and exact answers and gold standard snippets. The questions come in four categories (i.e. factoid, list, yes/no and summary). Factoid and list questions are usually answered by a word or phrase (multiple word/phrases for list questions) making them amendable for extractive answer span detection type exact question answering for which general purpose question answering data sets are available such as SQUAD [13]. Snippets matching their corresponding exact answer(s) are selected for the bio-medical question answering labeled set generation. For about 30% of the factoid/list questions no snippet can be aligned with their corresponding ideal answers. We analyzed those cases and were able to recover additional 152 questions after manual inspection for synonyms and transliterations to include in our labeled data set. The labeled data set is split into 85%/15% training/testing data sets of size 9557 and 1809, respectively.

For yes/no answer classification, the ideal answer text of each BioASQ yes/no questions is used as the context and the exact answer (i.e. ‘yes’ or ‘no’) as label for binary classification. The ideal answers are cleaned up to remove the exact answer (yes or no) that sometimes occur at the beginning of the ideal answer. The labeled data is split into 85%/15% training/testing data sets of size 729 and 129, respectively. BioASQ yes/no questions are skewed towards yes answers where about 80% of the answers were yes.

For named entity recognition tests, we have used publicly available datasets used by Crichton et. al [14]. Four common biomedical entity types are considered, namely disease, drug/chemical, gene/protein and species. For each entity type one data set is selected for training/testing as summarized in Table I.

**TABLE I.**
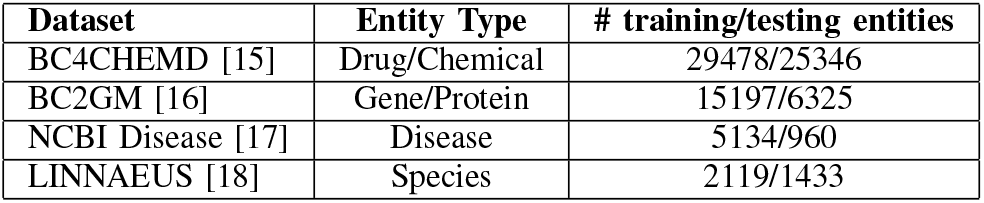
Bimedical named entity recognition data sets

| Dataset | Entity Type | # training/testing entities |
| --- | --- | --- |
| BC4CHEMD [15] | Drug/Chemical | 29478/25346 |
| BC2GM [16] | Gene/Protein | 15197/6325 |
| NCBI Disease [17] | Disease | 5134/960 |
| LINNAEUS [18] | Species | 2119/1433 |

### B. Effect of amount of pre-training on the Bio-ELECTRA performance

The effect of the increased number of training steps on the BioASQ question answering task is shown in Figure 1 on exact-match evaluation measure where the 95% confidence intervals are also shown, Even at 880K (or 440K in terms of ELECTRA Small++ pre=training with doubled batch size) training steps the performance of the Bio-ELECTRA is strong relative to BERT Base as shown in Table II. Similar to what is observed in general purpose downstream question answering tasks [1], [3], more pre-training improves downstream performance in biomedical question answering.

**Fig. 1.**
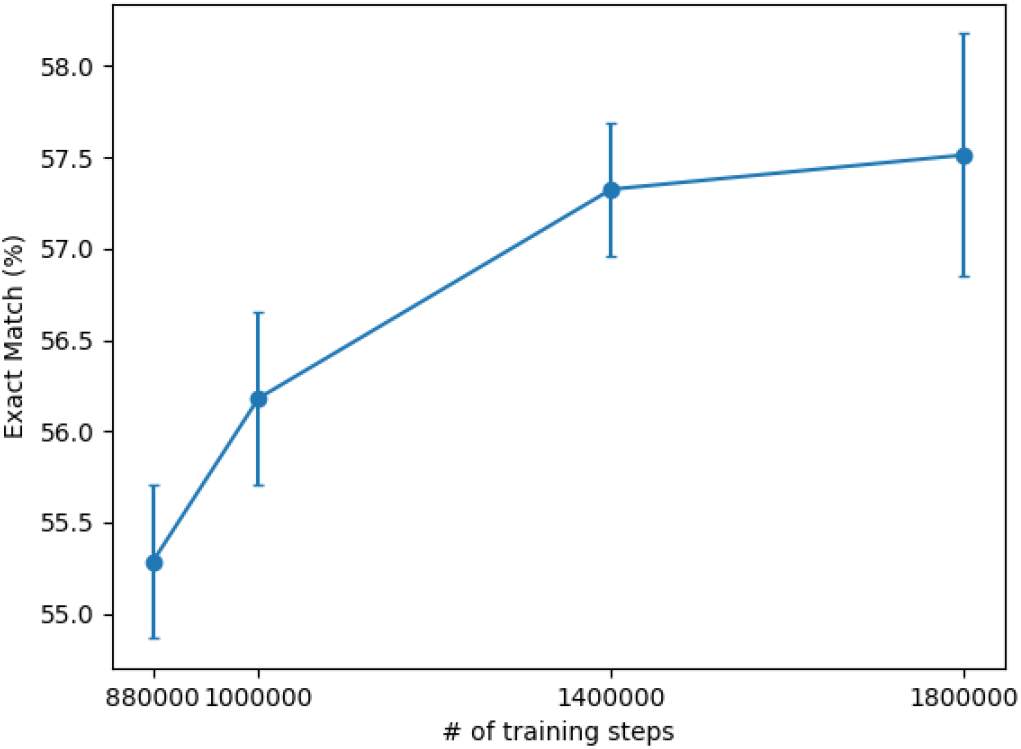
Change in the exact match performance for BioASQ question answering as a function of increased pre-training of Bio-ELECTRA

**TABLE II.**
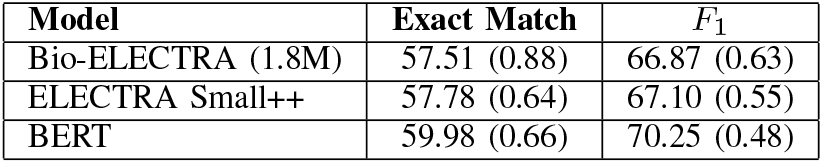
Biomedical Question Answering Test Results

### C. Experimental Results

The biomedical factoid/list question answering results are shown in Table II. We have used official SQUAD evaluation measures exact answer span match percentage and *F*_1_ measure. While BERT Base model had slightly better performance, taken into account their 8 times smaller size and 45 times less training time [3], the performance of both Bio-ELECTRA and ELECTRA Small++ models are impressive. With one fourth of the training of ELECTRA Small++, Bio-ELECTRA has nearly same performance as the ELECTRA Small++.

BioASQ yes/no question answer classification task results are shown in Table III. We have used the official BioASQ yes/no question evaluation measure of precision, recall and *F*_1_ applied on both yes and no questions separately. Here, Bio-ELECTRA outperforms BERT Base. The high standard deviations for Bio-ELECTRA and BERT Base are due to one random run in each case being stuck in a local minimum where the classifier always answers yes (since BioASQ yes/no questions are highly unbalanced towards yes answer (80% yes/20% no)).

**TABLE III.** Biomedical Yes/No Question Answer Classification Test Results

| Model | P (Yes) | R (Yes) | F<sub>1</sub> (Yes) | P (No) | R (No) | F<sub>1</sub> (No) |
| --- | --- | --- | --- | --- | --- | --- |
| Bio-ELECTRA (1.8M) | 87.99 (2.95) | 97.94 (1.35) | 92.66 (1.56) | 77.14 (26.47) | 46.92 (16.39) | 58.18 (19.91) |
| ELECTRA Small++ | 88.18 (0.71) | 94.31 (1.74) | 91.14 (1.00) | 69.92 (7.34) | 50.38 (3.19) | 58.40 (3.61) |
| BERT Base | 87.02 (2.57) | 95.49 (2.64) | 90.99 (1.00) | 65.15 (22.99) | 43.46 (15.20) | 51.71 (17.49) |

The test results for biomedical NER experiments are shown in Table IV. Similar to BioBERT [2], we have used precision, recall and *F*_1_ as evaluation measures. Here, large BERT Base language representation model shows, the largest benefit over smaller models at the cost 8 times longer inference time. Bio-ELECTRA was slightly better (in terms of mean F1 performance) than ELECTRA Small++ in three of the four NER entity types, while ELECTRA Small++ was slightly better than Bio-ELECTRA on Disease entity type.

**TABLE IV.** Biomedical Named Entity Recognition Test Results

| Type | Dataset | Metrics | ELECTRA Small++ | Bio-ELECTRA | BERT |
| --- | --- | --- | --- | --- | --- |
| Disease | NCBI disease | P | 76.14 (0.73) | 74.51 (0.57) | 85.43 (0.62) |
|  |  | R | 84.83 (0.71) | 81.94 (0.68) | 87.08 (0.76) |
|  |  | F<sub>1</sub> | 80.25 (0.60) | 78.04 (0.38) | 86.24 (0.55) |
| Drug/chem. | BC4CHEMD | P | 80.06 (0.40) | 80.66 (0.51) | 91.36 (0.13) |
|  |  | R | 81.88 (0.16) | 84.37 (0.58) | 89.46 (0.22) |
|  |  | F<sub>1</sub> | 80.96 (0.19) | 82.47 (0.14) | 90.40 (0.11) |
| Gene/protein | BC2GM | P | 67.56 (0.57) | 68.12 (0.48) | 83.95 (0.27) |
|  |  | R | 76.27 (0.39) | 76.00 (0.33) | 84.30 (0.31) |
|  |  | F<sub>1</sub> | 71.65 (0.38) | 71.84 (0.32) | 84.13 (0.23) |
| Species | LINNAEUS | P | 91.18 (0.59) | 91.35 (1.33) | 96.01 (0.31) |
|  |  | R | 93.64 (0.81) | 94.23 (0.67) | 93.90 (0.17) |
|  |  | F<sub>1</sub> | 92.39 (0.53) | 92.76 (0.87) | 94.94 (0.17) |

## IV. Conclusion

In this paper, we have shown that small domain-specific language representation models that make more efficient use of pre-training data can achieve comparable downstream performance on several biomedical text mining tasks to BERT Base with eight times more parameters. A domain-specific biomedical language representation model based on recently introduced ELECTRA architecture named Bio-ELECTRA is pre-trained on a consumer grade GPU with only 8GB memory.

While, Bio-ELECTRA performance is highly competitive to BERT Base for question answering and classification tasks, its performance lags behing BERT Base for NER tasks. To further improve the performance of Bio-ELECTRA, we are currently in the process of further pre-training with extended biomedical corpus of full papers from PMC open access initiative.

